# Parasitic helminth infections in humans modulate Trefoil Factor levels in a manner dependent on the species of parasite and age of the host

**DOI:** 10.1101/2021.06.10.447912

**Authors:** Babatunde Adewale, Christopher F. Pastore, Heather L. Rossi, Li-Yin Hung, Jeff Bethony, David Diemert, James Ayorinde Babatunde, De’Broski. R. Herbert

## Abstract

Helminth infections, including hookworms and Schistosomes, can cause severe disability and death. Infection management and control would benefit from identification of biomarkers for early detection and prognosis. While animal models suggest that Trefoil Factor Family proteins (TFF2 and TFF3) and interleukin-33 (IL-33) -driven type 2 immune responses are critical mediators of tissue repair and worm clearance in the context of hookworm infection, very little is known about how they are modulated in the context of human helminth infection. We measured TFF2, TFF3, and IL-33 levels in serum from patients in Brazil infected with Hookworm and/or Schistosomes, and compared them to endemic and non-endemic controls. TFF2 was specifically elevated by Hookworm infection, not Schistosoma or co-infection. This elevation was more strongly correlated with age than with worm burden. To determine if this might apply more broadly to other species or regions, we measured TFFs and cytokine levels in both the serum and urine of Nigerian school children infected with *S. haematobium*. We found that serum levels of TFF2 and 3 were reduced by infection, but urine cytokine levels were increased (IL-1β, TNFα, IL-13, and IL-10). Finally, to determine if TFF2 and 3 might have immunosuppressive effects, we treated stimulated or unstimulated PMBCs with recombinant human TFF2 or TFF3 and measured proinflammatory cytokine levels. We found that rhTFF2, but not rhTFF3, was able to suppress TNF alpha and IFN gamma release from stimulated human PMBCs. Taken together, these data support a role for TFF proteins in human helminth infection.

**Author Summary:** Billions of people are infected with parasitic helminths across the globe, especially in resource poor regions. These infections can result in severe developmental delay, disability, and death. Adequate management of helminth infection would benefit from the identification of host biomarkers in easily obtained samples (e.g. serum or urine) that correlate to infection state. Our goal was to determine if specific proteins involved in tissue repair and immune modulation are altered by infection with specific helminth species in Brazil (hookworm) or Nigeria (blood fluke). One of these proteins, Trefoil Factor 2 (TFF2), was elevated in the serum of hookworm infected individuals, and decreased in the serum of blood fluke infected children. In the blood fluke-infected children, there were also significant increases in pro-inflammatory cytokines in the urine, where the eggs burst from host tissue. Further, in laboratory experiments, Trefoil Factor 2 reduced the release of pro-inflammatory cytokines from human blood cells. This suggests that at high levels TFF2 may suppress inflammation and could serve as a biomarker for infection or treatment efficacy.

## Introduction

Billions of people are infected with parasitic helminths including cestode, nematode, and trematode species that are collectively responsible for millions of disability associated life years (DALYs) annually across the globe [1]. While considerable advances have been made in drug development and mass drug administration efforts can reduce worm burdens in endemic areas, the high prevalence of re-infection in humans living in endemic countries [2-4] suggests worms have evolved multiple ways to subvert and/or avoid host immunity. For instance, infection of children with hookworms including *Necator americanus* and *Ancylostoma duodenale* infections causes anemia and failure to thrive [5-8]. In adults, hookworm infections have been linked with a generalized immunosuppression that may lead to reduced vaccine efficacy [9, 10]. *N. americanus* experimental infections engage host immunoregulatory pathways driven by cytokines like interleukin 10 (IL-10) and transforming growth factor beta (TGF-β) [11], which could partially explain the ameliorative effect of hookworm infections in the context of autoinflammatory diseases, like celiac disease [12]. Whether such immunosuppression is due to host derived suppressive molecules and/or parasite derived factors released within the excretory secretory products remains unclear.

The immunoregulatory impact of parasitic helminths on their hosts is also a central feature of infections by blood flukes in the Schistosoma species. Human schistosomiasis, a neglected tropical disease that affects over 250 million people worldwide, results in severe morbidity, compromised childhood development, and an estimated 280,000 deaths annually [13]. Although praziquantel administration is an effective pharmacological treatment against adult Schistosomes [14], patients often present with elevated re-infection rates [15, 16], which again points to the ability of Schistosomes to modulate the immunological landscape of their hosts during chronic parasitism. Indeed, humans infected with *S. haematobium* can mount prototypical Type 2 responses associated with reduced damage and parasite clearance [17], but also show reduced tendency toward allergic responses [18-20], implying that ongoing infection downmodulates the inflammatory status of the host. Further, the generation of long-lasting immunotherapies to combat Schistosoma infections is greatly limited by our poor understanding of how helminth-induced inflammation is regulated. A greater understanding of how different worm species module their hosts is certainly needed in order to address three unresolved issues: 1) does parasite burden corelate with or predict the degree of immunomodulation, 2) do humans living in distinct endemic areas mount similar responses, 3) how does tissue injury caused by parasitic infection elicit host immune and tissue repair responses?

Amongst all of the known tissue repair mechanisms operating at the mucosal interface, Trefoil factor family proteins (TFF1, TFF2, and TFF3) remain one of the most poorly understood. TFF proteins are small secreted glycoproteins produced by goblet cells under both homeostatic and injury-induced conditions [21-23]. Trefoils are named after their evolutionarily conserved cloverleaf shaped “P” or trefoil-domain, which imparts functional resistance to proteolysis [24]. TFF2 and TFF3 are the predominant TFFs produced in the colon of humans and most mammals, but only bear ∼20% amino acid conservation [25]. TFFs 1-3 are diagnostic for GI tissue injury responses in human mucosa, e.g., TFF expression marks the ulcer-associated cell lineage (UACL), which defines cells located at the regenerative border of GI ulcers [26, 27]. Although it is known that prophylactic administration of rTFF2 and rTFF3 into the GI lumen of rodents with injury induced inflammation leads to suppression of disease [28-30], the lack of protective efficacy for TFF3 enema administration in a clinical trial of humans with Ulcerative Colitis [31] has led to controversy regarding the necessary context and utility for using TFFs in the regulation of GI inflammatory disease. Our work has implicated TFF’s as important immunoregulatory molecules in the context of GI parasite infection [32-35]. Importantly we found that TFF2 can suppress proinflammatory cytokine production in the context of *T. gondii* infection [34] and the genetic deficiency in TFF3 led to increased release of Type 1 inflammatory cytokine INFγ [32].

The overarching goal of this study was to address the idea that the chronic injurious nature of hookworm or Schistosoma infection in different parts of the world would associate with changes in TFF2 or 3. We also sought to determine if parasite intensity might correlate with the levels of TFFs. Here, we find that hookworm infection in a Brazilian cohort preferentially elevated TFF2 levels, even when compared to co-infection with Schistosomes. In contrast to our expectation, we found that older age, rather than egg burden had a stronger positive correlation with TFF2 levels. In agreement with this, children in Nigeria with *S. haematobium* infection exhibited lower levels of serum TFF2 and TFF3, which corresponded with higher levels of cytokines in the urine, including the type 2 cytokine IL-13, the proinflammatory cytokines TNFα and IL-1β, and the regulatory cytokine IL-10. Interestingly, exposure of human PBMC from normal subjects not living in endemic areas shows that TFF2, but not TFF3 had suppressive effects on the ability of these cells to undergo PHA induced proinflammatory cytokine production (TNFα and IFN-γ). Taken together, this work indicates that TFF levels are modulated in body fluids of infected individuals and may have a role in promoting immunoregulation that occurs in such individuals.

## Methods

### IRB Approval and Recruitment of Samples from Human Subjects

The George Washington University IRB approved the project (IRB# 100310), which was effective August 22, 2011. Samples for this study were recruited from this ongoing study from 2013-2014. The study was conducted in an area endemic for hookworm in Americaninhas, Novo Oriente, Northeastern Minas Gerais, Brazil. All communication about the study, including the written consent form was conducted in Portuguese. The Brazilian Ministry of Health (MOH) and local health officials from each village were in charge of the on-site medical supervision. These officials routinely supply and administer anti-helminthics, and are proficient and skilled at drawing blood in a rural setting, due to continuous surveillance studies run by the MOH. Prior to obtaining written consent, subjects were informed of the study during a village meeting, when members of the local departmental health institutes provided an explanation about the aim, execution plan, and methodologies of the study. At this meeting the villagers were able to ask questions and offer their opinions, and efforts were made to ensure that the village residents understood, including their right to refuse participation in the study. After this meeting, written consent was obtained from all adult subjects, from the parents or guardians of minor subjects, and written assent was obtained from the minor subjects. Exclusion criteria included full time school or work attendance outside of the endemic area, a positive pregnancy test, or hemoglobin < 80g/L.

The George Washington University IRB also approved an additional protocol (IRB# 190988) to recruit healthy adult subjects (18-50 years old) to provide control serum samples via word of mouth and advertisement (e.g. newspaper, online, Research Match). These subjects were also informed of the purpose of the study via in person meeting with a member of the study team in a private room, who also answered any questions the subject might have and stress that participation is voluntary. Subjects also receive a written description of the study and provided their signature on an informed consent form. It was made clear that they could withdraw their sample at any time, and the written document provided a means to contact the research team if they chose to revoke consent. Blood collection was conducted by a trained individual and the volume of collection did not exceed 50 mL.

The Nigerian Institute of Medical Research IRB approved the project (No. IRB/18/042), which was effective April 4, 2018 – March 10, 2019, when patient information and samples were recruited. Social approval for the study was obtained from the Medical Officer in-charge of Health in the Local Government as well as the approval of the Education Secretary of the Borgu Local Government Education Authority. Details of the procedure were explained to all participants during the social mobilization stage. The written informed consent of parents was obtained with the assent of the children to participate in the exercise. Participation was voluntary and the assent of each child was obtained before sample collection. The study was conducted in accordance with the tenets of Helsinki Declaration of 1964 as amended in 2013 and guidelines of Good Clinical Practice.

### Patient Demographics and Samples

#### Brazilian Cohort

Patients were recruited from hookworm (*N. americanus)* endemic areas for this study, consented to provide serum and fecal samples (to diagnose infection and assess egg burden). TFF2, TFF3, and IL-33 were measured from serum samples by ELISA. Initial analysis indicated that sexes did not differ. Further, endemic and non-endemic controls did not differ in their levels of any analyte, and were pooled for statistical comparison with patient levels. The first experiment assessed co-infection state, which is common in the region, and included patients infected with either Schistosoma (n = 11, 6 female), Hookworm (n = 20, 9 female), or both (n = 16, 8 female), and were compared to both uninfected endemic (n = 25, 13 female) and non-endemic (n = 20, 10 female) controls (data in Fig.1A-C). This cohort ranged 5-67 years old (27 mean ± 18 SD years). We also collected from a larger cohort of hookworm-infected individuals that included 69 females (3-72 years old, 29 mean ± 21 SD years) and 97 males (4-70 years old, 27 mean ± 20 SD years) (data in Fig.1D-F).

**Figure 1.**
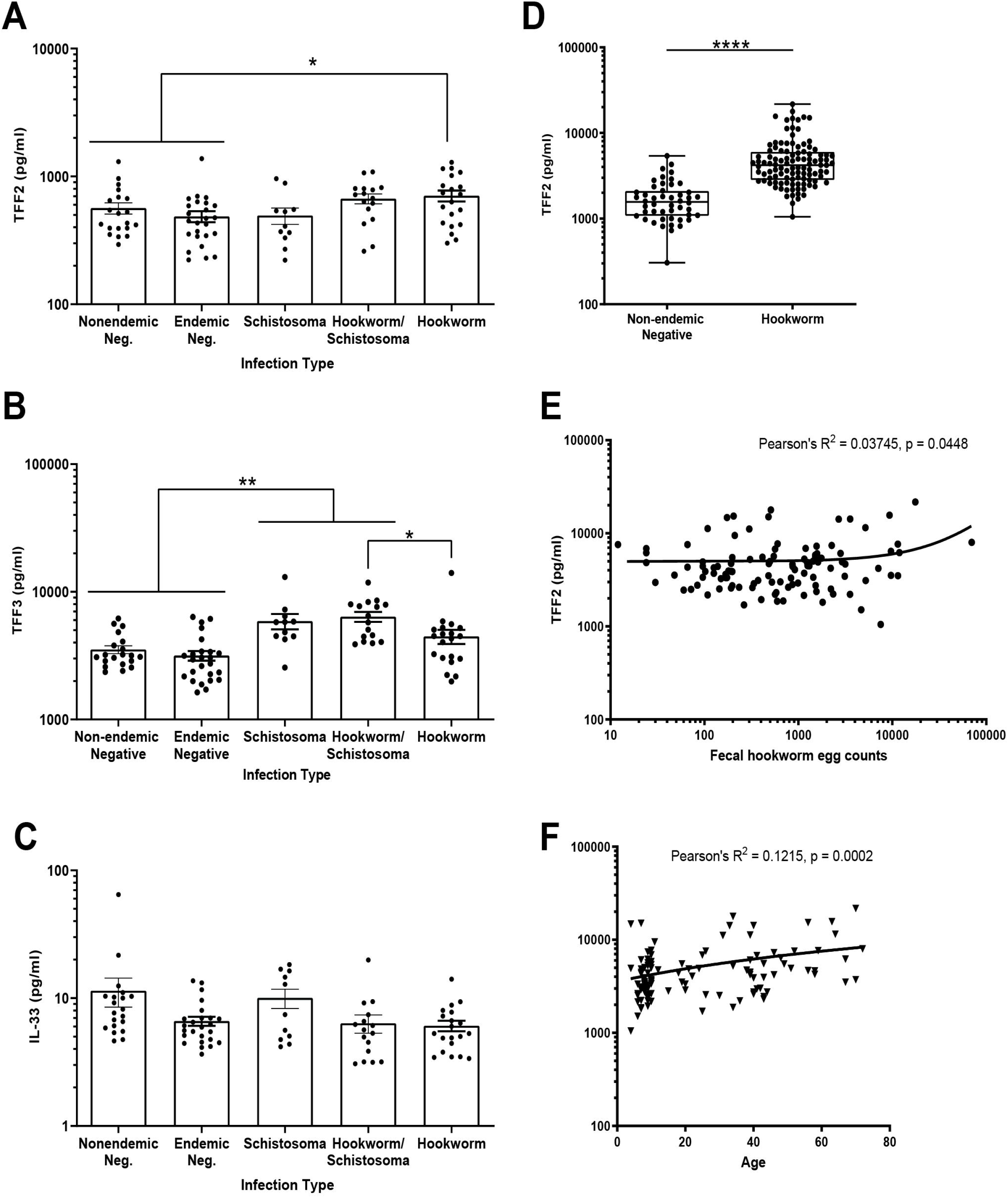
TFF2 and 3 levels are modulated differently by hookworm versus Schistosoma helminth species and TFF2 levels are age-dependent in individuals from Brazil. (A) Levels of TFF2 or (B) TFF3 in serum samples from Brazilian patients with either Schistosoma (n = 11), Hookworm (n = 20), or both (n = 16) versus uninfected controls (n = 25 endemic and n = 20 non-endemic). (C) Levels of TFF2 in serum sample from hookworm-infected patients (n = 108) and non-endemic negative controls (n = 48). (D) Linear relationship between patient fecal egg burden and serum TFF2 levels. (E) Linear relationship between patient age and serum TFF2 levels in hookworm-infected patients. * p<0.05, ** p<0.01, **** p<0.0001

#### Nigerian Cohort

Children aged 6-17 years undergoing routine health screening through their school provided blood, urine and feces, and were divided into infected or uninfected groups based on the presence of eggs from the blook fluke *S. haematobium* in urine samples. Fecal samples were used to determine if any control individuals (no *S. haematobium* eggs in urine) also had hookworm infection and these were excluded from analysis. There were no *S. haematobium* infected children co-infected with hookworm. Trefoil factors and cytokines (IL-33, IL-1b, IL-13, IL-17A, IL-22, IL-10, IFNγ and TNFα) were measured from serum and urine samples by ELISA. Dipstick uranalysis was performed by pipetting drops of urine onto Multisitx 10SG reagent strips from Siemens (Tarrytown, NY), to assess for signs of damage to the urinary tract such as elevated protein and presence of blood in the urine, and other features (specific gravity, pH, ketones, glucose, nitrite, urobilinogen, and leukocyte content).

In total, 78 (30 female) children, aged 6-14 years, provided samples. Preliminary laboratory analysis done at the Nigerian Institute of Medical Research was included prior to shipment to the University of Pennsylvania, where ELISA measurements were performed. After shipment, some samples lacked sufficient volume for testing. Of the serum samples tested, 51 came from children without *S. haematobium* eggs in their urine (23 were female, 17 had hookworm eggs in feces), 16 came from infected children with *S. haematobium* eggs in their urine (2 were female). Of the urine samples tested, 48 came from children without *S. haematobium* eggs in their urine (21 were female, 14 had hookworm eggs in feces), 17 came from infected children with *S. haematobium* eggs in their urine (23 were female). Only hookworm uninfected controls (n = 34) were compared to *S. haematobium* infected samples (n = 16-17).

### Culture and Treatment of Human Peripheral Blood Mononuclear Cells

Human Peripheral Blood Mononuclear Cells (PBMCs) were obtained from four anonymous donors and generously provided by Dr. Douglas Nixon. Cells were plated 2.5 × 10^5^ per well in RPMI buffer (Gibco, Amarillo, TX) containing 5-10% human non-autologous plasma (Gemini Bio, West Sacramento, CA) and maintained in 37°C incubation with 5% CO_2_. Some wells were pre-treated with human recombinant TFF2 or 3 overnight (25 ng/µL, US Biologicals, Salem, MA), before being stimulated with phytohemaglutinin (PHA, 50µg, Thermofisher, Waltham, MA), for 24 hours. Unstimulated controls were run simultaneously, both with or without exogenous TFFs, to confirm a lack of direct effect on cytokine release. Supernatants were collected and IFNγ and TNFα were measured by ELISA at the end of stimulation. Each treatment point is the average of two duplicate wells.

### Enzyme Linked Immuno-Sorbent Assays (ELISAs)

Commercially available ELISAs were used to measure levels of human TFF2, TFF3, IL-33, IL-1β, IL-13, IL-17A, IL-22, IL-10, IFNγ and TNFα in human serum and urine samples, and for IFNγ and TNFα, media from control or stimulated human PMBCs, according to the manufacturer’s instructions. Human TFF2, TFF3, and IL-33 ELISA kits were from R&D Systems (Minneapolis, MN). Human IL-1β, IL-10, IL-22, IFN-γ, and TNF-⍰ ELISA kits were from Biolegend (San Diego, CA). Human IL-13 and IL-17A ELISA kits were from eBioscience (San Diego, CA). Because the Brazilian cohort required multiple plates to process the samples, standard curves and positive and negative plate to plate quality controls were run on each plate to ensure consistency across plates. All standard curves produced R squared values ranging from 0.978-0.993, and the quality controls were less than 2 standard deviations from the average across plates, indicating consistent performance across plates.

### Statistical Analyses

Graph Pad Prism (v9) was used to graph and statistically analyze the data. One statistical outlier, defined as greater than 2 standard deviations above the group mean, was excluded from the endemic control TFF2 serum data set in Fig.1D for the Brazilian cohort. For two group comparisons, Welch’s t-tests were used. For comparisons between three groups, one-way ANOVA was used, with post-hoc Tukey’s test. Pearson’s correlations were used to determine the contribution of egg count or age to TFF2 levels in hookworm infected patients. For culture data, which does not conform to normality, non-parametric comparison of three groups was performed using the Kruskal-Wallis test, with post-hoc Dunn’s multiple comparisons test. Data in box and whisker plots are expressed as median and the interquartile range, while data in bar graphs are expressed as mean and standard error of the mean.

## Results

### Helminth Infections Differentially modulate TFFs and Hookworm Single Infection Increases TFF2 Age-dependently in the Brazilian Cohort

We have previously shown critical roles for TFF2, TFF3 and the alarmin cytokine IL-33 in protective immune responses in a murine model of hookworm infection [32, 33, 36], which needs to be confirmed in human infection. We first addressed this in patient samples from Brazil. Serum levels of TFF2, TFF3, and IL-33 did not differ between endemic and non-endemic controls (Fig. 1A-C), so these were pooled for comparison against patient samples. Co-infection of helminths is quite common in Brazil, particularly between hookworm (*N. americanus*) and *Schistosoma mansoni*, but there do not seem to be common genetic factors associated with their regulation within a single host, suggesting they may have different effects on immune responses [37]. Therefore, we sought to determine what effect co-infection of hookworm and Schistosoma, or Schistosoma infection alone might have on TFF2, TFF3, and IL-33 levels in serum. There was a significant effect of infection on serum levels of TFF2 (F_3, 88_ = 3.404, p = 0.0211) and TFF3 (F_3, 88_ = 11.88, p<0.0001), but not on IL-33 levels (F_3, 88_ = 1.246, p = 0.2980) (Fig. 1A-C). Hookworm infection alone was associated with significantly elevated TFF2 in serum as compared to controls, while Schistosoma-only or coinfection were intermediate and not significantly different from controls (Fig. 1A). In contrast, TFF3 levels in serum from Hookworm-infected individuals was no different from pooled controls and significantly lower than Hookworm/Schistosoma co-infection, while Schistosoma infection and co-infection had significantly greater TFF3 than controls (Fig. 1B). Given the trend toward increased TFF2 associated with Hookworm-only infection, we tested additional samples against endemic controls, and found a significant increase (t = 8.861, df = 135.9, p<0.0001, Fig. 1D). Taken together, this indicates that TFF levels are differentially regulated by different types of helminth infection. Specifically, TFF2 elevation is specifically associated with Hookworm infection while TFF3 elevation is associated with Schistosoma infection in a mostly adult population.

Given that our cohort from Brazil may have variable levels of worm burden and have a wide age-range (2-72 years), we sought to assess whether these factors correlate positively or negatively with TFF2 levels in hookworm-infected patients. There was a small but significant positive correlation for both fecal egg count (R^2^ = 0.03745, n = 108, p = 0.0448, Fig. 1E) and age (R^2^ = 0.1215, n = 108, p = 0.0002, Fig. 1F). This indicates that age, rather than worm burden, is more positively correlated with TFF2 levels in hookworm-infected individuals.

### TFF Serum Levels are Decreased with *S. haematobium* Infection in Nigerian School Children

Helminths impact a large proportion of the globe and a wide range of age groups. Nigeria is another region where different helminth infections are common, including the blood fluke *Schistosoma haematobium*, which is more prevalent among Nigerian school children than *S. mansoni*[38]. Given the differences we observed in the Brazilian cohort between hookworm and Schistosoma infections, we also sought to characterize TFF2, TFF3, and IL-33 levels in a pediatric cohort from Nigeria including children infected with *S. haematobium*. We found that both TFF2 (t = 5.896, df = 30.04) and TFF3 (t = 8.317, df = 42.94) were significantly decreased in serum samples from children infected with *S. haematobium* (Fig. 2A, B), but not IL-33 (Fig. 2C). Although this species of blood fluke infects the urinary tract and causes damage during infection[39], we did not find a similar pattern of significance in urine samples from infected versus uninfected children (Fig. 2D-F), indicating that urine cannot substitute for serum in terms of immune-related responses in this infection. TFF2 levels are much higher in urine than serum levels for the whole cohort, and TFF3 levels also seem to be higher in urine than serum in some cases. This may reflect a high degree of clearance of these factors. As indicated previously (Fig. 1D), lower age is associated with lower TFF2 levels, indicating the low measured values of the Nigerian children versus the mostly adult Brazilian cohort. Taken together, these data indicate that helminth species differentially affect the levels of TFF2 and TFF3 in an age-dependent manner.

**Figure 2.**
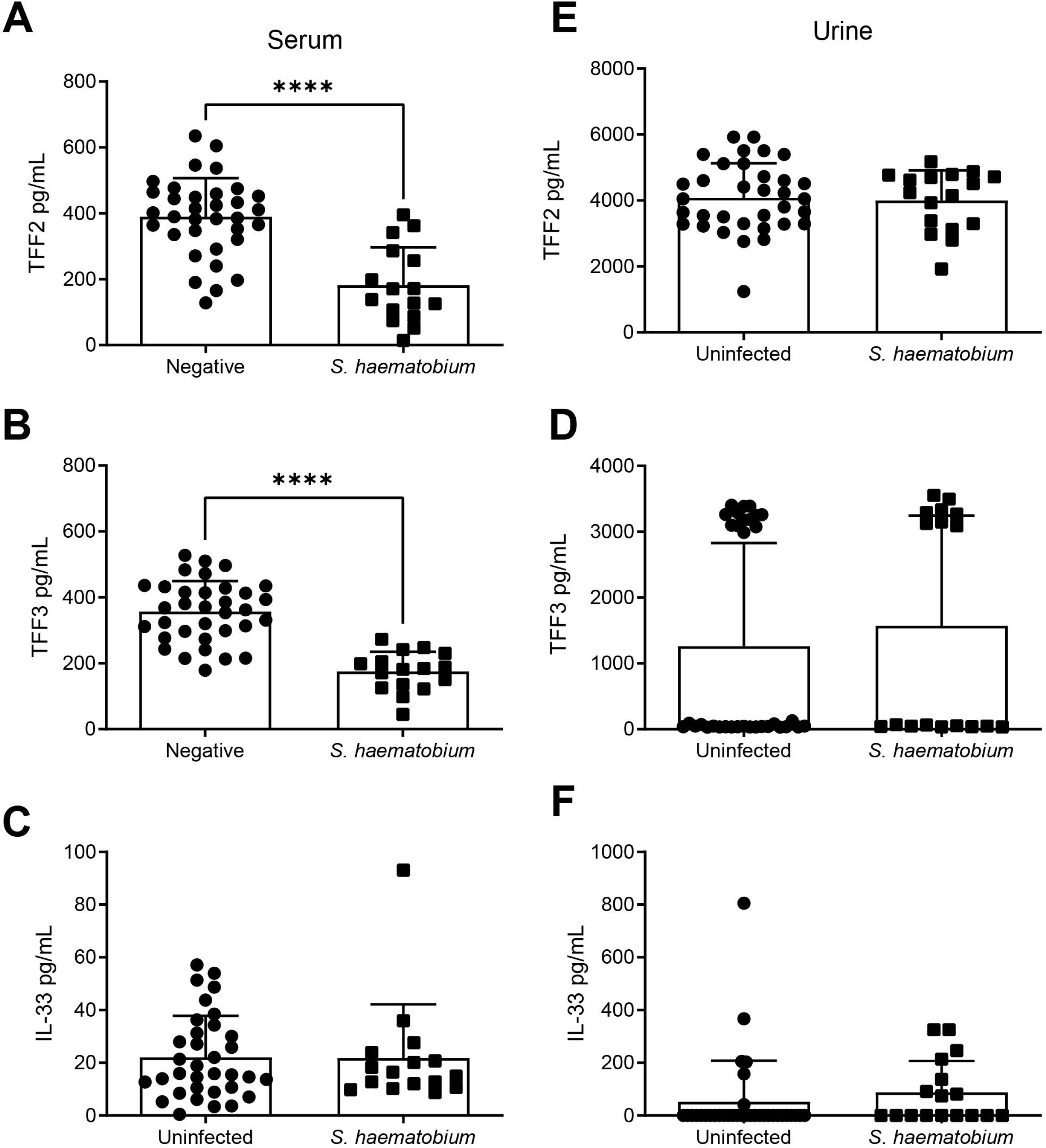
TFFs are decreased in serum but not urine samples from Nigerian children infected with *S. haematobium*. (A, B) Levels of TFF2 or (C, D) TFF3 in serum (A, C) and urine (B, D) samples from Nigerian school children infected with *S. haematobium* (n = 16-17) versus uninfected controls (n = 34). **** p<0.0001.

### *S. haematobium* Infection is Associated with Increases in Cytokines in Urine but not Serum

We further evaluated consequences of *S. haematobium* infection on cytokine levels in serum and urine, where serum may reflect systemic inflammatory responses, while urine may reveal damage-associated signals as well as site-directed healing. Indeed, urine samples from children with *S. haematobium* infection had significantly elevated protein (0.07353 vs 1.029, t = 4.650, df = 16.74, p = 0.0002) and blood (0.1029 vs 1.765, t = 4.884, df = 17.54, p = 0.0001) as compared to uninfected controls, but across other general measurements of the urine samples (e.g., glucose, pH, specific gravity, etc.) did not differ significantly (data not shown). Helminth infection is associated with alarmin cytokine signaling, and clearance is broadly associated with Type 2 immune responses [40]. In both compartments, we measured IL-1β, which is in the same family as IL-33 [41], cytokines associated with proinflammatory immune responses and enhanced tissue damage (IFNγ, TNFα) [42], with Type 2 immune responses (IL-13) [40], and other regulatory or healing processes (IL-17A, IL-22, IL-10) [43, 44]. As with TFF levels, serum and urine samples revealed differing patterns of increased cytokine levels in infected versus uninfected children. Serum levels were not significantly different for any cytokine tested (Fig. 3A-D and SFig. 1), although IL-13 (t = 1.974, df = 17.85, p = 0.0640) and IL-10 (t = 1.878, df = 15.77, p = 0.0790) trended towards significance. In urine, there was a significant increase in IL-13 (t = 3.735, df = 19.78, p = 0.0013), IL-1β (t = 2.547, df = 16.2, p = 0.0214), IL-10 (t = 3.290, df =16.08, p = 0.0046), and TNFα (t = 2.488, df = 19.48, p = 0.0221) in infected versus uninfected children (Fig. 4 E-H). We found no difference between infected versus uninfected for levels of IFNγ, IL-17A, or IL-22 in urine (Sfig.1). Taken together, these data indicate that significant elevations in inflammatory cytokines (IL-13, IL-1β, TNF α) and the regulatory cytokine IL-10 are detected in at the active site of damage in the urinary tract (urine), rather than more systemically in the serum.

**Figure 3.**
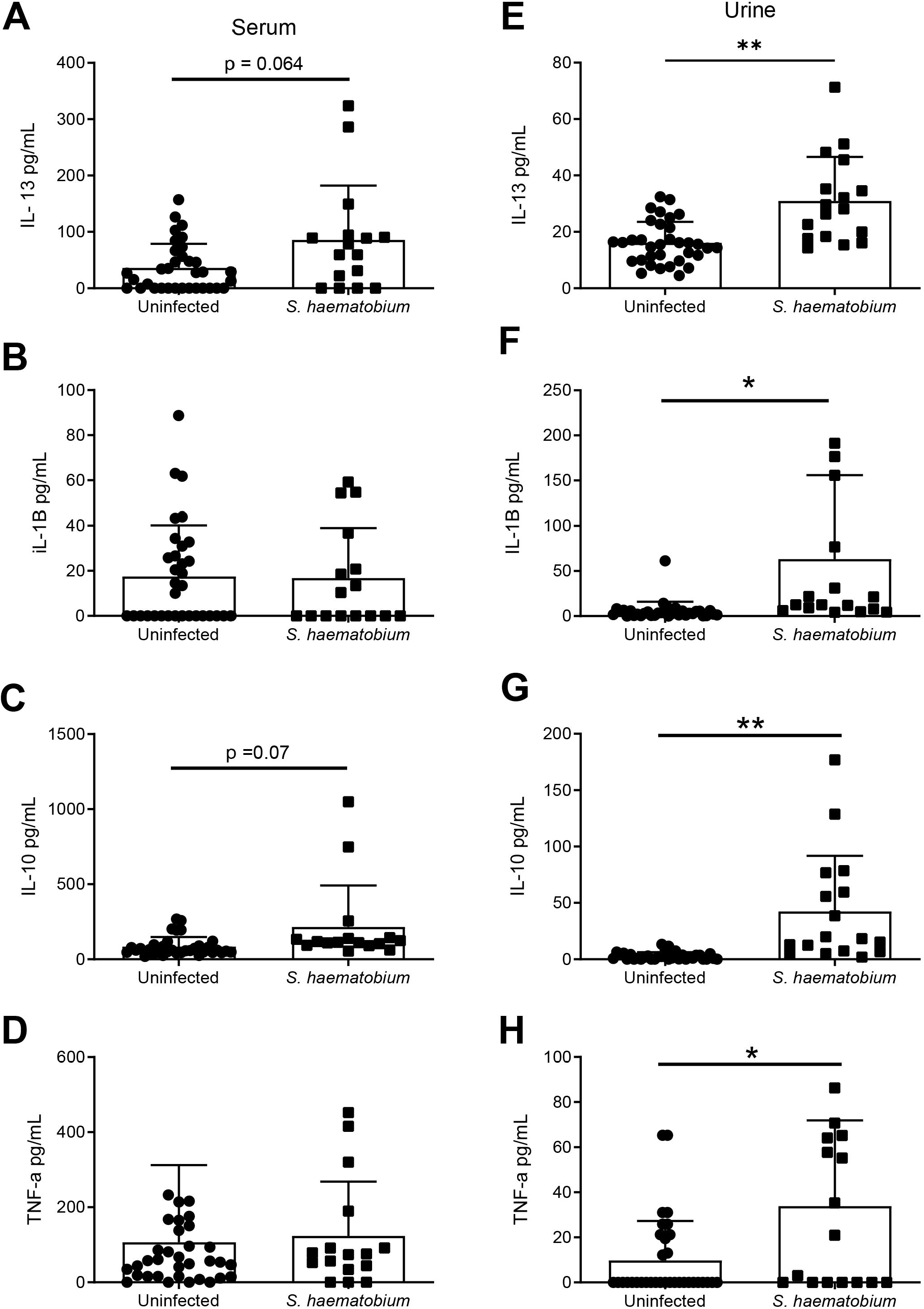
Cytokine levels are significantly elevated in urine, but not serum samples of Nigerian children infected with *S. haematobium*. (A-D) Serum levels and (E-H) urine levels of (A, E) IL-13, (B, F) IL-1b, (C, G) IL-10, and (D, H) were measured infected (n = 16-17) and uninfected (n = 34) children, and only urine samples from infected children were significantly higher. * p<0.05, ** p<0.01

**Figure 4.**
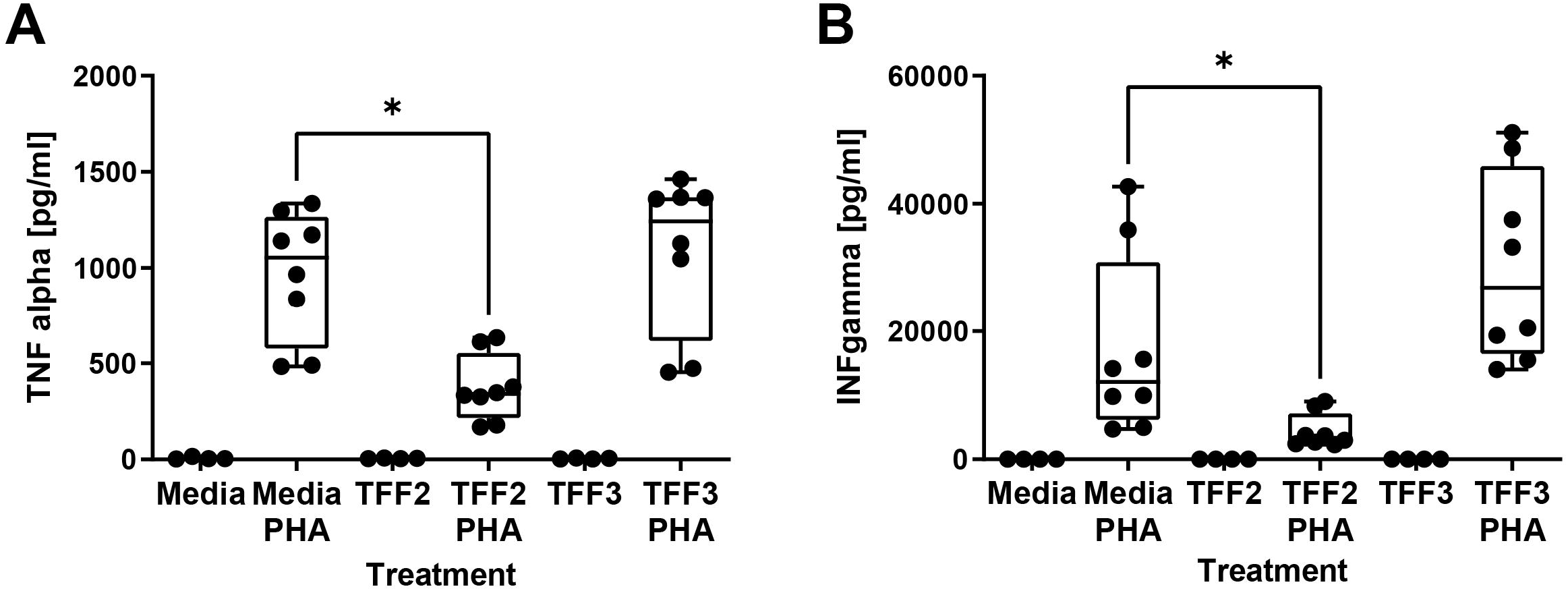
rhTFF2, but not rhTFF3 reduces PHA-evoked TNF alpha and INF gamma produced by human PMBCs in culture. (A) TNF alpha levels or (B) INF gamma levels produced by cultured PMBCs (2.5 × 10^5^ cells/well) following treatment with media alone, media with rhTFF2 or rhTFF3 (25 ng/µL each, n = 4 wells), PHA (50µg) alone or PHA with rhTFF2 or rhTFF3 (n = 8 wells). Pairwise comparisons as indicated: * p<0.05

### TFF2, but not TFF3, Suppresses Pro-Inflammatory Cytokine Release

Because TFFs are altered by helminth infection and known to modulate immune responses, we sought to determine what effect TFFs might have on proinflammatory responses. We cultured human PBMCs and stimulated them with phytohemaglutinin (PHA), both alone and in the presence of recombinant human TFF2 or TFF3. PHA mimics a highly inflammatory state and produces a significant increase in both interferon gamma (IFN-y) and tumor necrosis factor alpha (TNF-a) production vs media alone, or with TFFs alone (Fig. 4). This response was significantly repressed by co-treatment with rh-TFF2, but not rh-TFF3 (Fig. 4), indicating that elevated TFF2 may contribute to suppressed inflammatory responses in the context of hookworm infection.

## Discussion

Our previous work in mouse models of helminth infection strongly suggests that trefoil factors contribute to protective immune responses and parasite clearance [32-35]. To further support the translational potential of TFFs as a biomarker of infection or therapeutic target, we evaluated the levels of TFF2 and TFF3 for the first time in human helminth infection in geographically distinct regions. In Brazil, we found that human TFFs are modulated by helminth infection in a species and age-dependent manner. TFF2 was specifically elevated by hookworm infection and increased more strongly with age than with parasite burden. In contrast, infection with the blood fluke *S. haematobium* decreased TFF2 and TFF3 levels in serum of infected Nigerian children. This reduced level of TFFs was accompanied by increased levels of cytokines in the urine of infected children, including proinflammatory IL-1b and TNFα, the type-2 cytokine IL-13, and the inhibitory IL-10. Exogenous TFF2 was associated with suppressed TNFα and IFNγ production by stimulated human PMBCs, suggesting that elevated TFF2 in the context of hookworm infection could reduce these pro-inflammatory cytokine levels through direct or indirect mechanisms *in vivo*. Given that TFF2 is low in younger populations and further reduced in *S. haematobium* infected children, this could be why TNFα is high in urine and by proxy the infection site of the bladder and urinary tract, where *S. haematobium* eggs burst through tissue into the urine. These findings bring us closer to understanding the mechanisms involved in human helminth infection. This information is necessary for uncovering diagnostic biomarkers to evaluate treatment efficacy and to develop better strategies to control infection.

It is generally accepted that tissue damage caused by the invading larval stages of various worm species drives prototypical Type 2 responses [45-49], which contribute to worm clearance and tissue repair [40]. These type 2 responses are initiated by the release of alarmin cytokines like IL-25, IL-33 or, thymic stromal lymphopoietin (TSLP) from damaged tissue [45-49]. We were unable to detect any significant differences in IL-33 levels in either human cohort, suggesting that this cytokine is not a good biomarker of infection. This finding does not necessarily negate a potential role for IL-33 in human immune responses to helminths, as it may be relevant at the infection site at specific times during the course of infection, which may not be adequately sampled by minimally-invasive methods.

Alarmin cytokines are key signals to activate tissue resident type 2 innate lymphocytes (ILC2s) and CD4+ TH2 cells, leading to their proliferation and release of both type 2 cytokines (IL-13 and IL-5) [48]. We were able to detect significant elevation in IL-13 associated with *S. haematobium* infection, particularly at the site of damage, supporting the idea that type 2 immune responses occur in the context of human infection. This may suggest that IL-13 in urine could serve as a non-invasive biomarker to track treatment efficacy. Additionally, *S. haematobium* is difficult to model in experimental animals, but some features can be recapitulated in a mouse model system [50]. Our findings in human infection largely concur with earlier findings in the mouse model, specifically chronically elevated IL-13 and TNFα in the bladder tissue of egg-injected mice and only transient changes in IL-17 [50]. Some of the regulatory cytokine findings did differ between human data and the mouse model. Specifically, IL-10 elevation, but not TGFβ was associated with human infection, whereas the opposite was true in the mouse model [50]. This supports the translational relevance of the mouse model system, as well as a potential avenue to uncover mechanisms that could be targeted to boost immunity in humans.

These type 2 responses are thought to both augment clearance mechanisms against helminth colonization and also act to initiate tissue repair [40]. Host tissue repair would be advantageous for the parasite, as helminth survival depends on the host’s survival. TFF proteins are also critical for mucosal tissue repair [51], and may act upstream or in parallel with type 2 immune responses. Although our work in mouse models of hookworm infection suggests that both TFF2 and TFF3 contribute to worm clearance, tissue repair, and the suppression of proinflammatory cytokines [32-35], we found that TFF2 is specifically increased in the context of human hookworm infection and can reduce secretion of proinflammatory cytokine by human PMBCs. In the mouse hookworm infection model *N. brasiliensis*, TFF2 is required for induction of IL-33, early Type 2 cytokine responses, and contributes to clearance of worms from the gut[35]. We found that high TFF2 in human hookworm infection was not associated with higher IL-33 levels. This may suggest a species difference that could be worth examining further. The lack of IL-33 induction in humans may explain why the infection persists. Further exploration as to why TFF2 in mice can induce IL-33, but may not do so as strongly in humans could help identify mechanisms where human immunity could be boosted to improve worm clearance.

In contrast to hookworm infection, *S. hematobium* infection in children was associated with a decrease in both TFF2 and TFF3 in the serum, but not the urine. The reason why TFFs modulation is detectable in serum, but not urine, while cytokine differences were seen only in urine and not serum is not entirely clear. It could point to different responses of the host to the adult form found in vasculature versus the eggs encysted in the bladder wall and released into the urine [52]. This is the first time TFFs have been examined in the context of *S. hematobium* infection, but there is some link in the literature connecting TFFs to bladder [53-55]. Of note for our findings, cats with idiopathic cystitis, a painful condition characterized by hypercontractivity and frequent urination, have reduced levels of TFF2 in bladder biopsies compared to healthy controls [54]. It may be useful to determine if TFF levels correlate with measurements of bladder dysfunction or other aspect of *S. hematobium* infection in the future. Given the age and species specific-effects on TFF2 levels in serum of the Brazilian cohort, it will be necessary to expand these studies in children to additional age groups in the future.

## Conclusion

It is critically important that we find ways to identify helminth infected individuals using easily obtained samples and inexpensive techniques that can be implemented in endemic regions. One of these approaches is the identification of disease biomarkers. Our findings provide an initial step toward determining if TFFs or other immune mediators might serve as biomarkers for specific helminth infections. Disease biomarkers could be used to track the efficacy of drug treatments for parasite clearance and to focus expensive mass drug administration efforts on the most vulnerable populations, like children at risk for developmental delay, which will minimize risk of generating drug resistant parasites, particularly Schistosomes [15, 56]. While the use of molecular diagnostic approaches, including parasite-derived microRNAs shows promise [57, 58], these have not been developed for *S. haematobium*. Our work suggests that evaluation of tissue reparative molecules that significantly change during active infection could present a viable approach for serological-based analyses in resource limited settings. It is likely that a combinatorial approach that uses both detection of parasite-derived and host-derived molecules would allow a highly specific approach to identify individuals with distinct parasite species and those who are likely to have a blunted response to vaccination against viral and bacterial pathogens.

## Acknowledgements

We thank Taylor Oniskey for her technical assistance regarding the Brazilian data associated with this project. This work was supported by NIH grants (U01-AI125940, R21-AI144572 awarded to DRH) and the Burroughs Wellcome Fund.

## Figure Legends

**Supplemental Figure 1.**
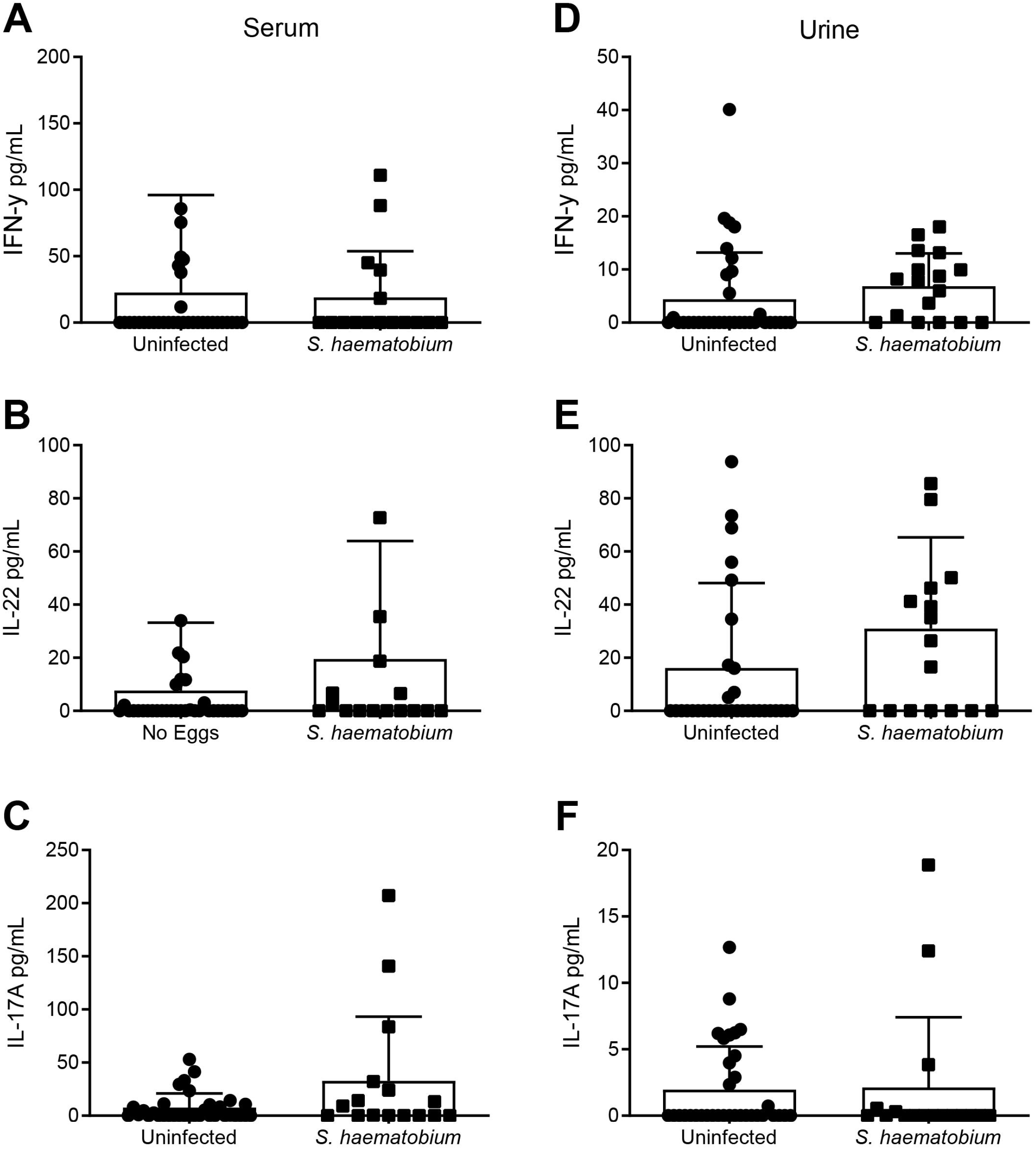
Cytokines (IFNγ, IL-22, and IL-17A) not changed in serum or urine samples of Nigerian children infected with *S. haematobium*. (A-C) Serum and (D-F) Urine samples were assessed for (A, D) IFNγ, (B, E) IL-22, and (C, F) IL-17A levels, and do not significantly differ between infected (n = 16-17) and uninfected (n = 34) children.

